# Divergent effects of *Wolbachia* on host temperature preference

**DOI:** 10.1101/2020.06.11.146977

**Authors:** Michael T.J. Hague, Chelsey N. Caldwell, Brandon S. Cooper

## Abstract

Heritable symbionts can modify a range of ecologically important host traits, including behavior. About half of all insect species are infected with maternally transmitted *Wolbachia*, a bacterial endosymbiont known to alter host reproduction, nutrient acquisition, and virus susceptibility. Here, we broadly test the hypothesis that *Wolbachia* modify host behavior by assessing the effects of eight different *Wolbachia* strains on the temperature preference of six *Drosophila melanogaster*-subgroup species. Four of the seven host genotypes infected with A-group *Wolbachia* strains (*w*Ri in *D. simulans, w*Ha in *D. simulans, w*Sh in *D. sechellia*, and *w*Tei in *D. teissieri*) prefer significantly cooler temperatures relative to uninfected genotypes. Contrastingly, when infected with divergent B-group *w*Mau, *D. mauritiana* prefer a warmer temperature. For most strains, changes to host temperature preference do not alter *Wolbachia* titer. However, males infected with *w*Sh and *w*Tei experience an increase in titer when shifted to a cooler temperature for 24 hours, suggesting that *Wolbachia*-induced changes to host behavior may promote bacterial replication and influence *Wolbachia* transmission rates. Modifications to host temperature preference likely influence host thermoregulation, and understanding the fitness consequences of these effects is crucial for predicting evolutionary outcomes of host-symbiont interactions, including how *Wolbachia* spread to become common.

## INTRODUCTION

Heritable symbionts have diverse ecological effects on their hosts. In insects, microbial symbionts influence host reproduction (e.g., cytoplasmic incompatibility; Hoffmann and Turelli 1997; Werren et al. 2008), acquisition of nutrients (Baumann 2005; Moran et al. 2008; Douglas 2009), tolerance of extreme temperatures (Brumin et al. 2011; Mueller et al. 2011), and susceptibility to viruses (Hedges et al. 2008; Teixeira et al. 2008). Much less is known about symbionts’ effects on host behavior and their ecological consequences (Feldhaar 2011; Goodacre and Martin 2012; Bi and Wang 2019; Hosokawa and Fukatsu 2020). On the one hand, symbionts may induce behavioral changes that promote the spread of infection through host populations. Because symbiotic relationships can span a continuum from mutualism to parasitism, behavioral modifications that promote infection spread may not necessarily benefit hosts (Hentschel et al. 2000; Werren et al. 2008). Parasites, for example, can induce behaviors that are detrimental or lethal to hosts, such as altering host locomotor behavior to increase the probability of parasite transmission (Lefevre and Thomas 2008; Poulin 2010; van Houte et al. 2013; Heil 2016; Vale et al. 2018; Weinersmith 2019). On the other hand, infected hosts may modify their own behavior in ways that mitigate negative aspects of the infection (Hart 1988; Poulin 2010; De Roode and Lefèvre 2012; Curtis 2014), such as a “behavioral chill” thermoregulatory response in which hosts seek cool temperatures to increase their survival probability (Fedorka et al. 2016). These behavioral effects represent an important component of how symbionts impact host fitness, which ultimately dictates the evolutionary trajectory of host-symbiont interactions.

Maternally transmitted *Wolbachia* are the most common endosymbionts in nature, infecting the cells of about half of all insect species, as well as other arthropods (Werren et al. 2008; Zug and Hammerstein 2012; Weinert et al. 2015). *Wolbachia* and host phylogenies are often discordant (O’Neill et al. 1992; Raychoudhury et al. 2009; Gerth and Bleidorn 2017), and most *Drosophila* hosts have acquired *Wolbachia* via introgressive and/or horizontal transfer within only the last few thousand years (Conner et al. 2017; Turelli et al. 2018; Cooper et al. 2019). Maternal transmission occurs in the host germline, but *Wolbachia* also infect a variety of host somatic cells, including metabolic, digestive, and nervous system tissue (Dobson et al. 1999; Albertson et al. 2009; Pietri et al. 2016). The fitness consequences of *Wolbachia* in host tissues ultimately determine infection spread, and initial spread from low frequencies requires positive *Wolbachia* effects on host fitness (Caspari and Watson 1959; Hoffmann et al. 1990; Barton and Turelli 2011). Exactly how *Wolbachia* alter components of host fitness is poorly understood (Ross et al. 2019b), even though theoretical and population-level analyses indicate pervasive positive effects on host fitness (Hoffmann et al. 1990; Hoffmann and Turelli 1997; Kriesner and Hoffmann 2018; Turelli et al. 2018; Meany et al. 2019; Hague et al. 2020).

Symbionts are known to influence host thermal tolerance (Russell and Moran 2006; Dunbar et al. 2007; Mueller et al. 2011; Wernegreen 2012; Zhang et al. 2019), and two recent studies found that *D. melanogaster* lines infected with the *w*MelCS or *w*Mel *Wolbachia* strains tend to prefer cooler temperatures than uninfected genotypes (Arnold et al. 2019; Truitt et al. 2019). Modifications to host temperature preference (*T*_*p*_) have important implications for insects, because ectothermic performance and fitness explicitly depend on temperature (Angilletta et al. 2004; Martin and Huey 2008; Dillon et al. 2009; Garrity et al. 2010; Hoffmann and Sgro 2011; Condon et al. 2014; Rajpurohit and Schmidt 2016). Because *Wolbachia* infect most insects (Werren et al. 2008; Zug and Hammerstein 2012; Weinert et al. 2015), it is crucial to understand how infections alter host thermoregulation. Few past analyses of insect behavioral thermoregulation have accounted for *Wolbachia* (Dillon et al. 2009; Hoffmann 2010; Rajpurohit and Schmidt 2016).

Differences in *T*_*p*_ between infected and uninfected flies could arise from conflicting physiological requirements of *Wolbachia* and their hosts. *Wolbachia* titer in host bodies is sensitive to temperature fluctuations (López-Madrigal and Duarte 2019), such that exceedingly cool (<20°C) and warm (>25°C) temperatures reduce titer and the efficiency of maternal *Wolbachia* transmission (Clancy and Hoffmann 1998; Ulrich et al. 2016; Ross et al. 2017, 2019a; Foo et al. 2019; López-Madrigal and Duarte 2019; Hague et al. 2020). *Wolbachia*-induced changes to *T*_*p*_ could provide more favorable thermal conditions for bacteria replication in hosts. Alternatively, host-induced changes to *T*_*p*_ could represent a host behavioral response that reduces *Wolbachia* titer to mitigate negative aspects of infection (e.g., behavioral chill). It is still unknown whether observed changes to *T*_*p*_ increase or decrease *Wolbachia* titer (Arnold et al. 2019; Truitt et al. 2019).

Here, we broadly test for *Wolbachia* effects on host *T*_*p*_ across the *D. melanogaster* subgroup of flies. Our experiments include seven A-group *Wolbachia*-infected genotypes (*w*Ri in *D. simulans, w*Ha in *D. simulans, w*MelCS in *D. melanogaster, w*Mel in *D. melanogaster, w*Sh in *D. sechellia, w*Yak in *D. yakuba*, and *w*Tei in *D. teissieri*) and one B-group *Wolbachia*-infected genotype (*w*Mau in *D. mauritiana*), which diverged from A-group strains 6 to 46 million years ago (Meany et al. 2019). We find that hosts infected with four of the A-group *Wolbachia* strains (*w*Ri, *w*Ha, *w*Sh, and *w*Tei) prefer a significantly cooler *T*_*p*_ than uninfected flies of the same host genotype. In contrast, *D. mauritiana* infected with B-group *w*Mau have a significantly warmer *T*_*p*_. Notably, we find no evidence for *w*MelCS or *w*Mel effects on *T*_*p*_ of *D. melanogaster*, which may be due to host effects. Shifting infected adults from an intermediate temperature towards their *T*_*p*_ for 24 hours generally does not alter *Wolbachia* titer, but in a few instances, reductions in host *T*_*p*_ seem to promote *Wolbachia* replication. Our results motivate future work on the causes and consequences of *Wolbachia* effects on *T*_*p*_ and other host behaviors.

## METHODS

### Fly Lines

We broadly tested whether *Wolbachia* alter *T*_*p*_ of their *Drosophila* host species. We evaluated eight different *Wolbachia* strains infecting six different species in the *D. melanogaster* subgroup (Supplemental Table S1). For two of these host species, we tested multiple *Wolbachia*-infected genotypes: *w*MelCS- and *w*Mel-infected *D. melanogaster* and *w*Ri- and *w*Ha-infected *D. simulans*. With the exception of the *w*MelCS-*D. melanogaster* line (*Canton S Berkeley*), all our *Wolbachia*-infected genotypes were naturally sampled to form isofemale lines, such that single gravid females were collected from the field and placed individually in vials. *w*MelCS is only found at low frequency in global populations of *D. melanogaster* (Riegler et al. 2005; Nunes et al. 2008; Richardson et al. 2012), because the strain has been largely replaced by a recent sweep of *w*Mel in roughly the last 5,000 years (Riegler et al. 2005; Nunes et al. 2008; Richardson et al. 2012; Cooper et al. 2019). *w*MelCS was originally identified in the common laboratory strain *Canton Special* (Stern 1943; Stern and Schaeffer 1943; Riegler et al. 2005), and a sub-strain (*Canton S Berkeley*) was kindly provided to us by Michael Turelli. All lines were maintained on standard cornmeal media prior to experiments.

We generated *Wolbachia*-uninfected genotypes by treating each infected line with 0.03% tetracycline for four generations. In the fourth generation, we used PCR to confirm flies were cleared of *Wolbachia*. We amplified both the *Wolbachia* surface protein (*wsp*) and a second set of primers for the arthropod-specific *28S* rDNA that served as a positive control (Cooper et al. 2017; Meany et al. 2019). We then reconstituted the gut microbiome of the tetracycline-cleared flies by rearing them on food where infected males of the same genotype had fed and defecated for the prior 48 hours. Tetracycline-cleared flies were given at least three more generations before we conducted experiments to avoid detrimental effects of the antibiotic treatment on mitochondrial function (Ballard and Melvin 2007).

### Host Temperature Preference Assays

We assayed *T*_*p*_ of each genotype using a thermal gradient apparatus adapted from Matute et al. (2009) and Goda (2014). The rectangular thermal gradient comprised a 44 x 13 x 1 cm plate of aluminum with a removable Plexiglas lid (Supplemental Figure S1). The Plexiglas lid enclosed a 1 cm-high space above the aluminum plate that allows flies to move around on the thermal gradient. We created an air-tight seal between the aluminum plate and the Plexiglas lid using double-sided tape and C-clamps. To keep flies on the temperature-controlled aluminum plate and off the lid, the Plexiglas was coated with Fluon (BioQuip Products), a slick barrier that prevents insects from obtaining a foothold (Dierick 2007; Dankert et al. 2009). An LED light was placed above the apparatus to ensure light was evenly distributed across the entire thermal gradient.

All *T*_*p*_ assays were conducted in a cold storage room with a constant temperature of 5°C. A hot plate set to 90°C was placed under one end of the aluminum plate to create a thermal gradient. All experiments began once the apparatus achieved thermal stability after approximately 0.5 hours. The aluminum plate was subdivided into seven 10 x 6 cm sections (Supplemental Figure S1), and we recorded the temperature at the center of each section using a thermocouple (Digi-Sense Traceable®) prior to the start of each experiment. Temperature decreased linearly along the gradient (R^2^ = 0.92), ranging from a mean of 34°C at the warmest end (section 1) to 17°C at the coldest end (section 7). Mean temperatures at the center point of each section across all experiments are shown in Supplemental Figure S2 and Table S2.

The following protocol for our assay was adapted from previous experiments (Matute et al. 2009; Goda et al. 2014; Rajpurohit and Schmidt 2016; Truitt et al. 2019). Trial runs revealed that a sample size of 50-60 flies allowed flies to distribute across the gradient without overcrowding in preferred temperature ranges, which is consistent with prior studies (Matute et al. 2009; Truitt et al. 2019). Flies were reared in a 25°C incubator under a 12L:12D light cycle (Pericival Model I-36LL) on a standard food diet. For each genotype, we collected virgin flies as a batch and separated them into four treatment groups: uninfected females, infected females, uninfected males, and infected males. Flies were then maintained in groups of 60 in individual food vials and aged to 3-5 days old. We selected a single batch each day and ran all four treatment groups separately in a randomized order, such that all flies assayed on a given day were of the same batch and age. All experiments were run between 9am and 5pm. Before each run, we measured the temperature at the center of each section along the gradient, then transferred flies into the apparatus through a small hole located in the middle of the Plexiglas lid where the temperature averaged 22.7°C (Supplemental Table S2). Flies were allowed to choose their preferred temperatures along the gradient for 30 minutes (Matute et al. 2009; Goda et al. 2014; Arnold et al. 2019; Truitt et al. 2019). At the end of this period, we scored the numbers of flies in each section. A small subset of flies located on the Plexiglas lid were removed from the analysis (Goda et al. 2014). After each run, the thermal gradient was cleaned with ethanol and allowed to dry.

For each genotype, we analyzed the *T*_*p*_ data using generalized linear mixed models (GLMMs) and a Poisson error structure in R (R Core Team 2018) with the “glmer” function in the *lme4* package (Bates et al. 2015). We treated *T*_*p*_ of each fly as the dependent variable and included infection status, sex, an infection-by-sex interaction, fly age (3, 4, or 5 days), and the run order of each replicate over the course of the day (1^st^, 2^nd^, 3^rd^, or 4^th^) as fixed effects. The replicate ID of each run was included as a random effect. We then assessed the significance of fixed effects using an analysis of deviance with chi-squared tests. The *T*_*p*_ data for some genotypes more closely approximated a normal distribution (see Supplemental Table S3), so we conducted an analogous set of tests using linear mixed models (LMMs) with the “lmer” function in the *lme4* package. Here, we assessed significance of fixed effects using an ANOVA with Wald’s chi-squared tests. The LMMs produced qualitatively similar results (Supplemental Table S3) to the GLMMs (Table 1), so only results from the GLMMs are presented in the main text.

**Table 1.** Results from the GLMMs. Statistically significant fixed effects at *P* < 0.05 are marked in bold text with asterisks.

|  | wRi |  |  | wHa |  |  | wMelCS |  |  | wMel |  |  |
| --- | --- | --- | --- | --- | --- | --- | --- | --- | --- | --- | --- | --- |
| Explanatory variable | coefficient | $\chi^2$ | *P* value | coefficient | $\chi^2$ | *P* value | coefficient | $\chi^2$ | *P* value | coefficient | $\chi^2$ | *P* value |
| Infection Status | 0.069 | 6.158 | **0.013\*** | 0.063 | 6.148 | **0.013\*** | -0.017 | 1.285 | 0.257 | -0.004 | 0.031 | 0.860 |
| Sex | -0.060 | 4.341 | **0.037\*** | -0.070 | 6.907 | **0.009\*** | -0.007 | 0.224 | 0.636 | -0.046 | 3.490 | 0.062 |
| Age | -0.003 | 0.016 | 0.898 | -0.001 | 0.020 | 0.887 | -0.019 | 11.426 | **0.001\*** | 0.012 | 2.251 | 0.134 |
| Run Order | 0.001 | 0.013 | 0.909 | 0.009 | 1.002 | 0.317 | 0.011 | 4.914 | **0.027\*** | 0.005 | 0.366 | 0.545 |
| Infection * Sex | -0.013 | 0.099 | 0.754 | -0.016 | 0.186 | 0.666 | 0.002 | 0.005 | 0.943 | 0.021 | 0.368 | 0.544 |
| Sample Size | 1015 |  |  | 857 |  |  | 1727 |  |  | 1341 |  |  |

|  | wMau |  |  | wSh |  |  | wYak |  |  | wTei |  |  |
| --- | --- | --- | --- | --- | --- | --- | --- | --- | --- | --- | --- | --- |
| Explanatory variable | coefficient | $\chi^2$ | *P* value | coefficient | $\chi^2$ | *P* value | coefficient | $\chi^2$ | *P* value | coefficient | $\chi^2$ | *P* value |
| Infection Status | -0.091 | 7.540 | **0.006\*** | 0.042 | 4.531 | **0.033\*** | 0.002 | 0.007 | 0.936 | 0.038 | 8.264 | **0.004\*** |
| Sex | -0.052 | 3.175 | 0.075 | 0.009 | 0.238 | 0.626 | -0.010 | 0.179 | 0.673 | -0.017 | 1.694 | 0.193 |
| Age | -0.018 | 2.686 | 0.101 | 0.003 | 0.130 | 0.718 | 0.026 | 3.633 | 0.057 | -0.012 | 2.717 | 0.099 |
| Run Order | 0.020 | 3.968 | **0.046\*** | 0.011 | 3.187 | 0.074 | 0.009 | 1.320 | 0.251 | 0.001 | 0.086 | 0.770 |
| Infection * Sex | 0.036 | 0.680 | 0.409 | -0.029 | 1.151 | 0.283 | 0.011 | 0.108 | 0.742 | 0.011 | 0.366 | 0.545 |
| Sample Size | 818 |  |  | 1087 |  |  | 1056 |  |  | 2500 |  |  |

A preliminary analysis of the data revealed that flies seemed to form a bimodal distribution along the thermal gradient, with one cluster of flies located at the cold end of the gradient (section 7) where temperatures averaged about 17°C (Supplemental Figure S3). Given that 17°C generally falls below the average *T*_*p*_ of *Drosophila* species reported in previous experiments (Matute et al. 2009; Rajpurohit and Schmidt 2016; Arnold et al. 2019; Truitt et al. 2019), we hypothesized that flies were becoming immobilized in section 7 due to the cold temperature (see Dillon et al. 2009). A similar phenomenon has been identified for *Caenorhabditis elegans* in assays of *T*_*p*_—the movement speed of *C. elegans* is dependent on temperature, which can leave worms “trapped” in cold sections of a thermal gradient (Anderson et al. 2007). Thus, we removed the putatively immobilized flies in section 7 from each dataset and re-conducted our analyses. The analyses excluding section 7 are presented in the main text; however, including section 7 did not alter our findings of *Wolbachia* effects on *T*_*p*_ (Supplemental Table S4). We concluded that the dataset excluding immobilized flies represents a more biologically accurate measure of *T*_*p*_ for each genotype.

### Wolbachia *Sequencing and Phylogenomic Analysis*

We conducted a phylogenomic analysis to characterize the evolutionary relationships among *Wolbachia* strains included in this study. We also tested whether *Wolbachia* effects on host *T*_*p*_ exhibit phylogenetic signal (i.e. whether closely related *Wolbachia* strains have similar effects on *T*_*p*_). We obtained *Wolbachia* sequences from publicly available genome assemblies, which included *w*Ri (Klasson et al. 2009), *w*Ha (Ellegaard et al. 2013), *w*Mau (Meany et al. 2019), and *w*Yak and *w*Tei (Cooper et al. 2019). We also obtained raw Illumina reads for a *w*Sh-infected *D. sechellia* individual from a previously published dataset (Accession SRX3029362; Schrider et al. 2018). Importantly, two divergent *Wolbachia* strains may infect *D. sechellia*: A-group *w*Sh and B-group *w*Sn. In nature, *w*Sh singly infects some individuals, but it also occurs as a co-infection with *w*Sn (Rousset and Solignac 1995). We confirmed that our *D. sechellia* genotype (*PmuseumbananaI*) is singly infected with *w*Sh using qPCR primers described below, which can distinguish between A-group and B-group *Wolbachia*. Finally, we sequenced our *w*MelCS- and *w*Mel-infected *D. melanogaster* genotypes (*Canton S Berkeley* and *PC75*, respectively) to compare the sequence similarity of our variants of these strains to those used in the prior assay of *T*_*p*_ by Truitt et al. (2019) (Chrostek et al. 2013).

Tissue samples for genomic DNA were extracted using a DNeasy Blood & Tissue Kit (Qiagen). DNA quantity was tested on a Nanodrop (Implen) and total DNA was quantified by Qubit Fluorometric Quantitation (Invitrogen). DNA was cleaned using Agencourt AMPure XP beads (Beckman Coulter, Inc.) following manufacturers’ instructions, and eluted in 50 μl 1 × TE buffer for shearing. DNA was sheared using a Covaris E220 Focused Ultrasonicator (Covaris Inc.) to a target size of 400 bp. We prepared libraries using NEBNext® Ultra™ II DNA Library Prep with Sample Purification Beads (New England BioLabs). Final fragment sizes and concentrations were confirmed using a TapeStation 2200 system (Agilent). We indexed samples using NEBNext® Multiplex Oligos for Illumina® (Index Primers Set 3 & Index Primers Set 4), and 10 μL of each sample were shipped to Novogene (Sacramento, CA, USA) for sequencing using Illumina HiSeq 4000, generating paired-end 150 bp reads.

Reads were trimmed using Sickle version 1.33 (Joshi et al. 2011) and assembled using ABySS version 2.0.2 (Jackman et al. 2017). *K* values of 71, 81, and 91 were used, and scaffolds with the best nucleotide BLAST matches to known *Wolbachia* sequences with *E*-values less than 10^−10^ were extracted as the draft *Wolbachia* assemblies. For each genotype, we chose the assembly with the highest N50 and the fewest scaffolds (Supplemental Table S5). The *w*MelCS, *w*Mel, and *w*Sh genomes, along with the five previously published genomes were annotated using Prokka version 1.11, which identifies homologs to known bacterial genes (Seemann 2014). To avoid pseudogenes and paralogs, we only used genes present in a single copy with no alignment gaps in all of the genome sequences. Genes were identified as single copy if they uniquely matched a bacterial reference gene identified by Prokka. By requiring all homologs to have identical length in all of the *Wolbachia* genomes, we removed all loci with indels. A total of 214 genes totaling 181,488 bp met these criteria.

We also repeated this analysis to include the *w*MelCS and *w*Mel genomes used in Truitt et al. (2019). Here, we restricted our analysis to only *w*MelCS and *w*Mel *Wolbachia*, with the goal of comparing sequence similarity between the variants used in this study to those from Truitt et al. (2019). Given that many loci accumulate indels over time, the number of loci included in this analysis of *w*Mel-like *Wolbachia* was relatively high, with a total of 720 genes totaling 733,923 bp that met our criteria. Based on these 720 genes, our *w*MelCS variant infecting the *Canton S Berkeley* genotype was identical to the *w*MelCS variant used in Truitt et al. (2019). Our *w*Mel variant infecting the *PC75* genotype was also highly similar to *w*Mel used in Truitt et al. (2019), with only 0.000016% third-position pairwise differences (only 4 out of 244,641 third-codon positions).

We estimated a Bayesian phylogram of the 214 genes from the eight different *Wolbachia* strains using RevBayes 1.0.8 under the GTR + Γ model partitioned by codon position (Höhna et al. 2016). Four independent runs were performed for each phylogenetic tree we estimated, and in each instance, all four runs converged on the same topology. All nodes were supported with Bayesian posterior probabilities of 1.

We used the resulting phylogram to test whether *Wolbachia* effects on host *T*_*p*_ exhibit phylogenetic signal. For each genotype, we extracted the least square (LS) mean *T*_*p*_ for infected and uninfected flies from the GLMMs, and then used the change in LS mean *T*_*p*_ as a continuous character to calculate the maximum likelihood value of Pagel’s lambda (λ; Pagel 1999). A *λ*value of 1 is consistent with a model of character evolution that entirely agrees with the phylogeny (i.e. *Wolbachia* effects on *T*_*p*_ are strictly proportional to relatedness), whereas a *λ*value of 0 indicates that character evolution occurs independently of the phylogenetic relationships (Freckleton et al. 2002). We used a likelihood ratio test to compare our fitted value of λ to a model assuming no phylogenetic signal (λ = 0) using the “phylosig” function in the R package *phytools* (Revell 2012). We also employed a Monte Carlo-based method to generate 95% confidence intervals surrounding our λ estimate using 1,000 bootstrap replicates in the R package *pmc* (Boettiger et al. 2012).

### Host Temperature Shift Experiments

The thermal gradient assays revealed a main effect of *Wolbachia* on host *T*_*p*_ for several genotypes. Interestingly, the host genotypes infected with A-group *Wolbachia* strains (*w*Ri, *w*Ha, *w*Sh, and *w*Tei) preferred cooler temperatures than uninfected flies, whereas *D. mauritiana*—the only host genotype infected with B-group *Wolbachia* (*w*Mau)—preferred a relatively warmer temperature. Truitt et al. (2019) speculated that the altered *T*_*p*_ of infected flies represents a host-induced behavior to reduce *Wolbachia* titer and ameliorate the negative effects of infection. According to this hypothesis, shifting species infected with A-group *Wolbachia* to a cool temperature should reduce *Wolbachia* titer in host bodies (i.e. behavioral chill), whereas shifting *D. mauritiana* infected with *w*Mau to a warm temperature should reduce *Wolbachia* titer (i.e. behavioral fever). To test these predictions, we reared the five infected genotypes listed above at an intermediate temperature of 21.5°C. We separated female and male virgins, aged them to 3 days old at 21.5°C, and then shifted them to either a cold (18°C) or warm (25°C) incubator for 24 hours. Flies were separated by sex and maintained in groups of 40 in individual food vials throughout the course of the experiment. Following 24 hours of the cold/warm temperature treatment, flies were frozen in a −80°C freezer for subsequent analysis of *Wolbachia* titer. This design enabled us to determine if an increase or decrease in *T*_*p*_ of *Wolbachia*-infected adults alters *Wolbachia* titer.

We used qPCR to compare *Wolbachia* titer in flies shifted to 18°C vs. 25°C. Flies from each temperature treatment were homogenized together in groups of 10. The final samples included 6 biological replicates for each sex and temperature treatment. DNA was extracted using a DNeasy Blood & Tissue Kit (Qiagen). We used a Stratagene Mx3000P (Agilent Technologies) to amplify *Drosophila*- and *Wolbachia*-specific loci. In order to quantify titer of the five different *Wolbachia* strains, we utilized multiple combinations of *Drosophila* and *Wolbachia* qPCR primers (Supplemental Table S6). Efficiency curves were generated to confirm that each primer pair had adequate efficiency. All qPCR reactions were amplified using the following cycling conditions: 50°C for 2 minutes, 95°C for 2 minutes, and then 40 cycles of 95°C for 15 seconds, 58°C for 15 seconds, and 72°C for 1 minute. We used the average cycle threshold (*Ct*) value of three technical replicates for each sample. We estimated relative *Wolbachia* density as 2^Δ*Ct*^, where Δ*Ct* = *Ct*_*Host*_ − *Ct* _*Wolbachia*_ (Pfaffl 2001). We then used a Wilcoxon rank sum test to assess differences in titer between flies shifted to 18° and 25°C.

## RESULTS

### Wolbachia Infections Modify Host Temperature Preference

In total, we assayed the *T*_*p*_ of 12,944 flies in 379 replicates on the thermal gradient. Results from the eight different infected genotypes are summarized in Figure 1, Supplemental Figure S4, and Table S7. *Wolbachia* infection status had a significant main effect on host *T*_*p*_ for five genotypes: *w*Ri-infected *D. simulans* (χ^2^ = 6.158, *P* = 0.013), *w*Ha-infected *D. simulans* (χ^2^ = 6.148, *P* = 0.013), *w*Mau-infected *D. mauritiana* (χ^2^ = 7.540, *P* = 0.006), *w*Sh-infected *D. sechellia* (χ^2^ = 4.531, *P* = 0.033), and *w*Tei-infected *D. teissieri* (χ^2^ = 8.264, *P* = 0.004) (Table 1). These results were robust to whether the data were analyzed using GLMMs or LLMs (Supplemental Table S3) or to whether we included flies that chose the coldest section of the thermal gradient (section 7; Supplemental Table S4). Of the five *Wolbachia* strains with a significant effect on *T*_*p*_, all host genotypes infected with A-group *Wolbachia* preferred a cooler temperature than uninfected flies (Figure 2): *w*Ri-infected *D. simulans* preferred a LS mean temperature of 21.72°C ± 1.02 (± s.e.) compared to 23.12°C ± 1.02 for uninfected flies, *w*Ha-infected *D. simulans* preferred a LS mean of 23.56°C ± 1.01 compared to the uninfected mean of 24.89°C ± 1.01, *w*Sh-infected *D. sechellia* preferred a LS mean of 23.32°C ± 1.01 compared to the uninfected mean of 23.98°C ± 1.01, and *w*Tei-infected *D. teissieri* preferred a LS mean of 22.70°C ± 1.01 compared to the uninfected mean of 23.70°C ± 1.01. In contrast, *D. mauritana* infected with B-group *w*Mau preferred a warmer LS mean temperature of 21.15 °C ± 1.01 compared to the uninfected mean of 19.67°C ± 1.02.

**Figure 1.**
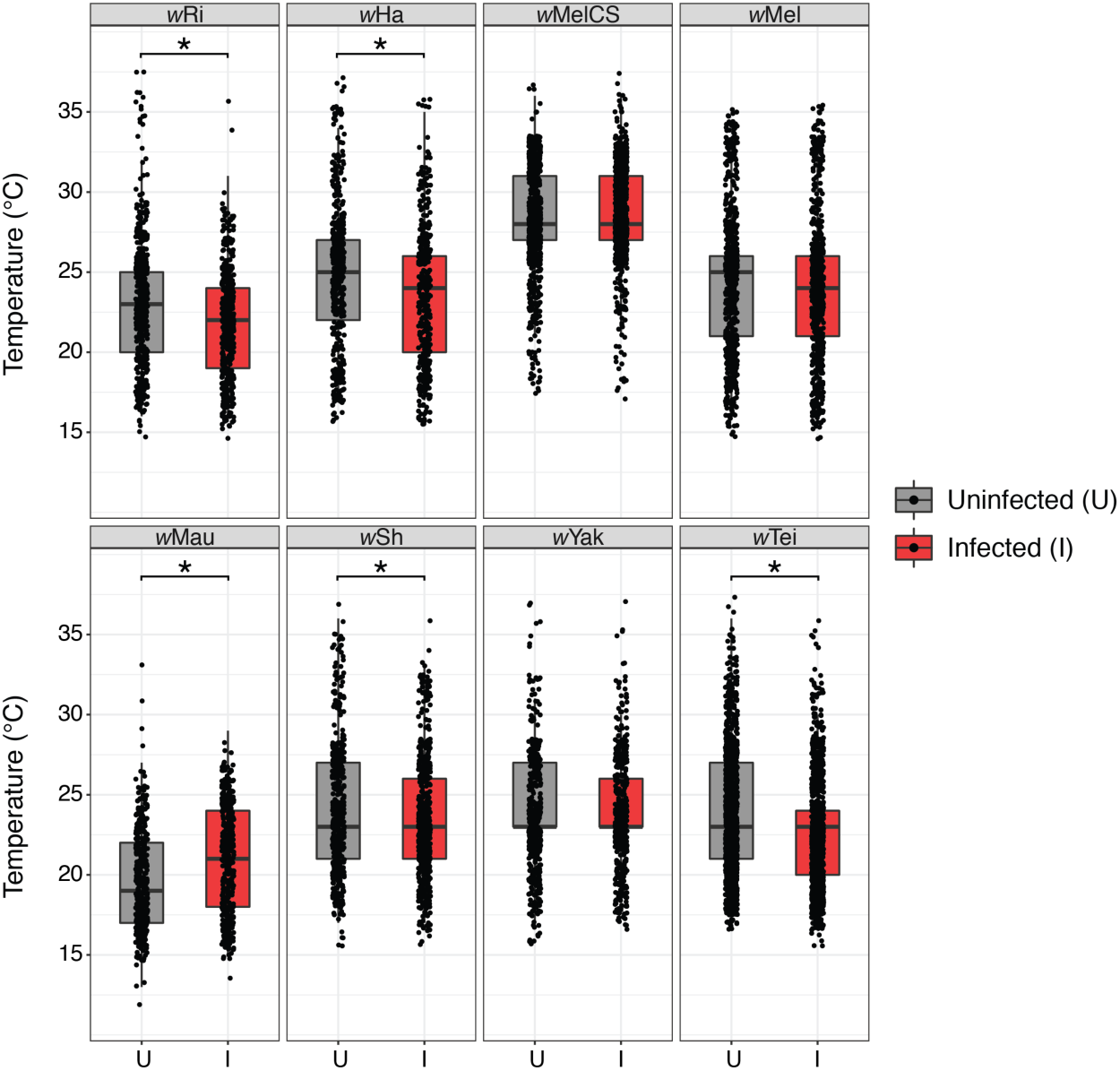
Box plots showing temperature preference (*T*_*p*_) for uninfected (U) and infected (I) flies of each genotype. Asterisks denote a significant main effect of *Wolbachia* infection on *T*_*p*_ from the GLMMs (Table 1). Individual points are jittered to show overlap. Similar plots including the coldest section of the thermal gradient (section 7) are shown in Supplemental Figure S3.

**Figure 2.**
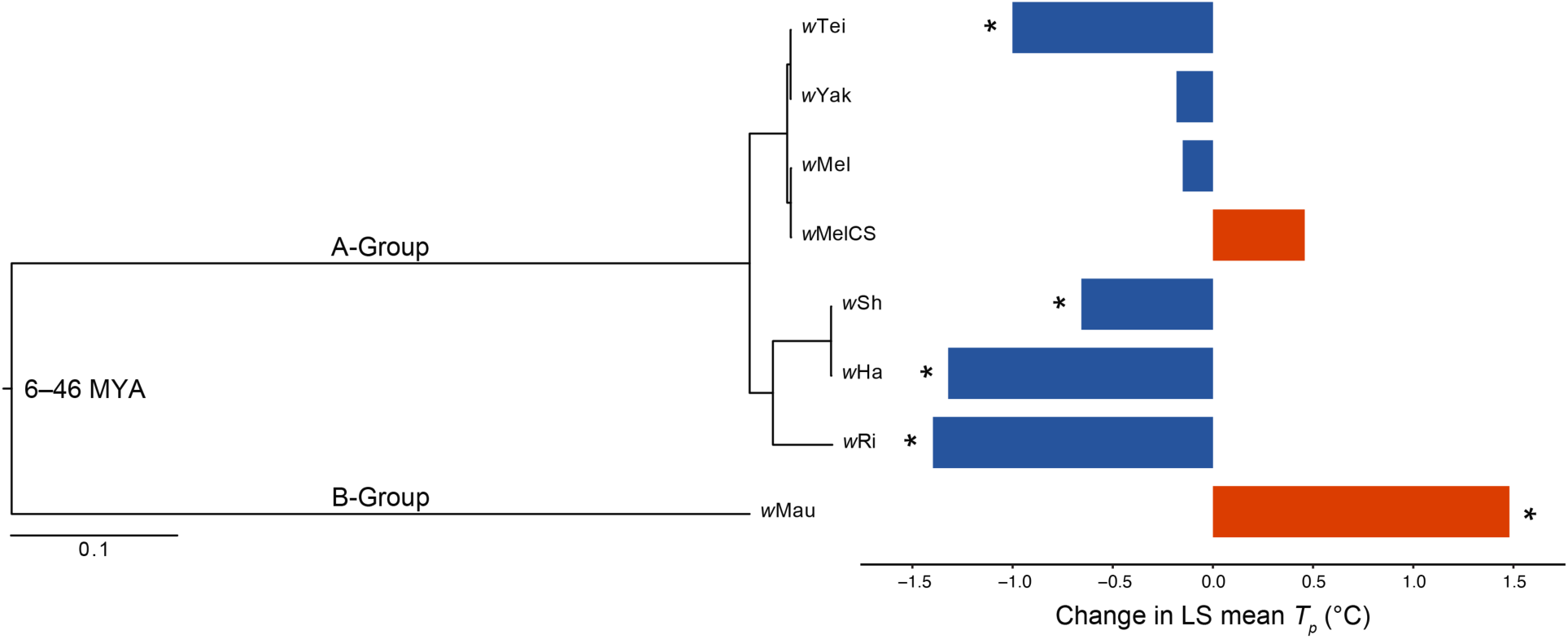
Estimated Bayesian phylogram for A- and B-group *Wolbachia* strains examined in this study. The phylogram was estimated with 214 single-copy genes of identical length in all of the genomes, spanning 181,488 bp. All nodes have Bayesian posterior probabilities of 1. To the right, the change in least square (LS) mean *T*_*p*_ between uninfected and infected flies is shown for each *Wolbachia* strain. LS means were generated from GLMMs (Table 1), and strains with a significant main effect on *T*_*p*_ are marked with asterisks. Divergence time estimates (MYA) for A- and B-group *Wolbachia* are from Meany et al. (2019).

In addition to *Wolbachia* infection status, we found other significant fixed effects on *T*_*p*_. Sex had a significant main effect on *T*_*p*_ for both the *w*Ri-infected *D. simulans* (χ^2^ = 4.341, *P* = 0.037) and *w*Ha-infected *D. simulans* (χ^2^ = 6.907, *P* = 0.009) (Table 1). For both of these *D. simulans* genotypes, males preferred cooler temperatures than females, regardless of infection status (Supplemental Figure S4). For the *w*Ri genotype, males preferred an LS mean temperature of 21.67°C ± 1.01 compared to the female mean of 23.17°C ± 1.01. For the *w*Ha genotype, males preferred an LS mean temperature of 23.29°C ± 1.01 compared to the female mean of 25.18°C ± 1.01. We also found a significant effect of fly age on *T*_*p*_ for *w*MelCS-infected *D. melanogaster* (χ^2^ = 11.426, *P* = 0.001), such that older flies tended to prefer cooler temperatures. Finally, we found that the run order each day had a significant effect on *T*_*p*_ for the *w*MelCS-*D. melanogaster* (χ^2^ = 4.914, *P* = 0.027) and the *w*Mau-*D. mauritiana* genotypes (χ^2^ = 3.968, *P* = 0.046). In both instances, flies assayed earlier in the day tended to prefer cooler temperatures. Interestingly, we only detected a main effect of *w*Mau on *D. mauritiana T*_*p*_ after accounting for run order—*w*Mau only had a marginal effect on *T*_*p*_ when we removed run order from the model (χ^2^ = 3.549, *P* = 0.060).

Next, we used the *Wolbachia* phylogram (Figure 2) to test whether *Wolbachia* effects on host *T*_*p*_ exhibit phylogenetic signal. Our fitted *λ*value was high (*λ*= 0.778 [0, 0.984]), but not significantly different from a model assuming no phylogenetic signal (likelihood ratio test, *P* = 0.203). The large confidence intervals associated with our *λ*estimate are likely due to the small number of strains included in the phylogram (*N* = 8). Further exploration of the distribution of *λ*estimates from the 1,000 bootstrap replicates showed a large number of near-zero values (Supplemental Figure S5). Small phylogenies are likely to generate near-zero *λ*values by random chance, not necessarily because the phylogeny is unimportant for trait evolution (Boettiger et al. 2012). To test whether larger phylogenies increase the accuracy of parameter estimation, we simulated trees with an increasing number of strains (*N* = 25, 50, and 100) and our *λ*estimate of 0.778 using the “sim.bdtree” and “sim.char” functions in the *geiger* R package (Harmon et al. 2008). See Supplemental Figure S5 for an extended description of the simulations. The simulated trees suggest that at least 50 strains are required to statistically distinguish *λ*≈ 0.8 from zero. The *N* = 25 tree had a fitted *λ*with extremely large confidence intervals (*λ*= 0.886 [0, 1]), whereas the *N* = 50 tree had a *λ*estimate that does not overlap with zero (*λ*= 0.860 [0.376, 0.977]). Together, these results suggest that *Wolbachia* effects on host *T*_*p*_ may exhibit phylogenetic signal, but our phylogeny is too small to distinguish statistically significant values of *λ*.

### *24-Hour Temperature Shifts Generally Do Not Alter* Wolbachia *Titer*

We tested whether shifting infected flies from an intermediate temperature towards their preferred temperature for 24 hours significantly alters *Wolbachia* titer. Results are summarized in Figure 3 and Supplemental Table S8. For *w*Ri-infected *D. simulans, Wolbachia* titer did not differ between the 24-hour cold and warm temperature treatments for females (*W* = 12, *P* = 1.000) or males (*W* = 19, *P* = 0.937). Similarly, for *w*Ha-infected *D. simulans*, titer did not differ between the temperature treatments for females (*W* = 13, *P* = 0.485) or males (*W* = 18, *P* = 1.000). We also observed no significant difference in titer between temperature treatments for *w*Mau-infected *D. mauritiana* females (*W* = 14, *P* = 0.589) or males (*W* = 14, *P* = 0.589). For *w*Sh-infected *D. sechellia*, we detected no difference in *Wolbachia* titer between females from each temperature treatment (*W* = 13, *P* = 0.485); however, we found that males significantly differed in titer between cold and warm treatments (*W* = 32, *P* = 0.026). Male *D. sechellia* shifted to 18°C had a higher median relative *Wolbachia* density (2^Δ*Ct*^ = 0.16) than males shifted to 25°C (2^Δ*Ct*^ = 0.11). This pattern is counter to the behavioral chill hypothesis proposed by Truitt et al. (2019), because *w*Sh-infected *D. sechellia* preferred a cooler temperature than uninfected flies, and shifting infected males to a cool temperature resulted in increased *Wolbachia* titer. We found the same pattern for *w*Tei-infected *D. teissieri*. While we detected no difference in *Wolbachia* titer between the treatments for females (*W* = 28, *P* = 0.132), males differed significantly in titer between the cold and warm treatments (*W* = 31, *P* = 0.041). As with *D. sechellia*, male *D. teissieri* shifted to 18°C had a higher median relative *Wolbachia* density (2^Δ*Ct*^ = 3.36) than males shifted to 25°C (2^Δ*Ct*^ = 2.98).

**Figure 3.**
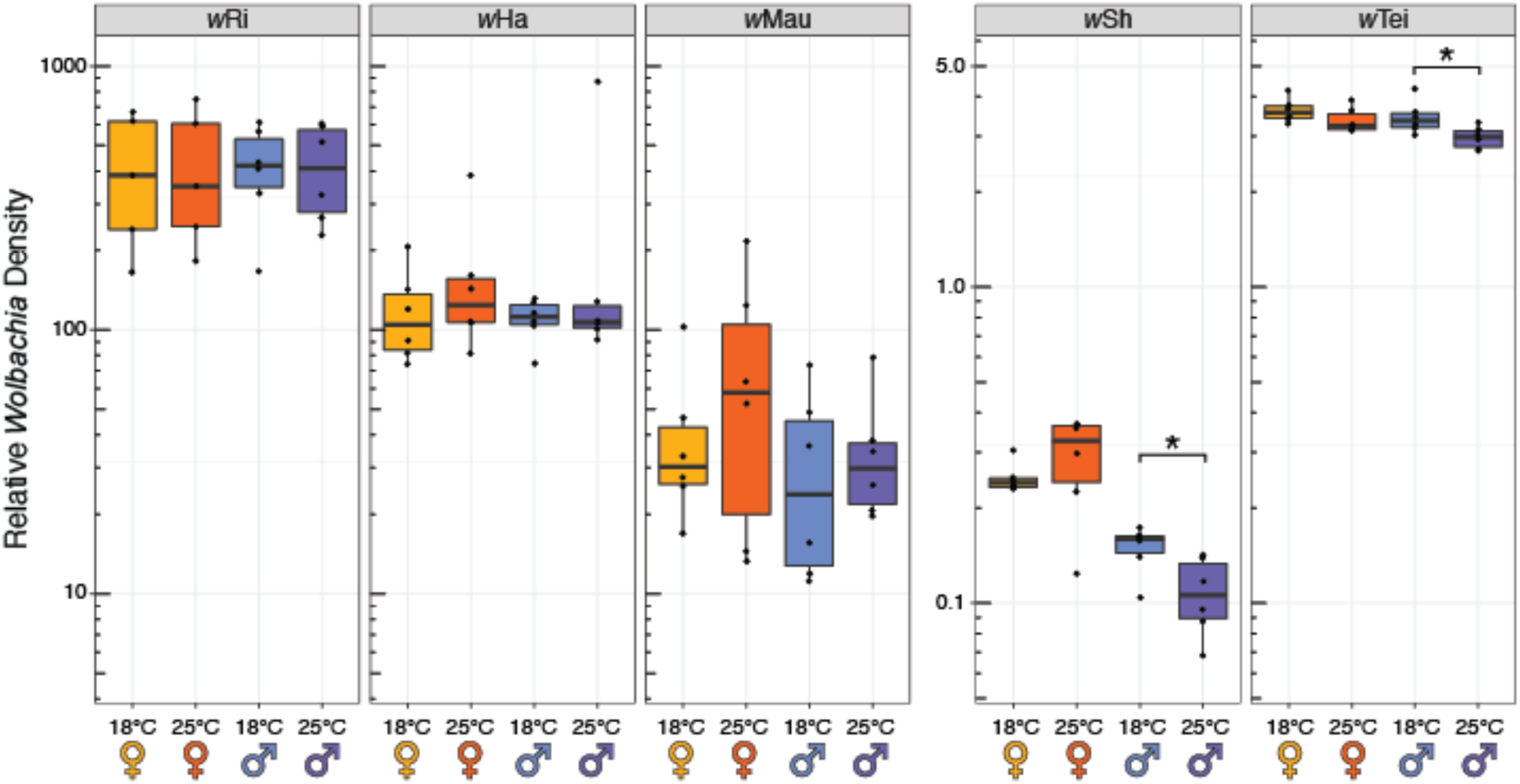
Boxplots of relative *Wolbachia* density from temperature shift experiments for the five *Wolbachia* strains showing main effects on host *T*_*p*_ (Table 1). Relative *Wolbachia* density is shown for virgin females (♀) and males (♂) shifted to cold (18°C) and warm (25°C) temperatures for 24 hours. Graphs are separated into strains with high titer (*w*Ri, *w*Ha, and *w*Mau) and low titer (*w*Sh and *w*Tei). Asterisks denote significant differences in titer between males shifted to 18°C and 25°C based on Wilcoxon rank sum tests.

## DISCUSSION

Our analyses suggest *Wolbachia* may generally influence host thermoregulatory behavior. Five of the eight *Wolbachia* strains we assayed had a significant effect on host *T*_*p*_: *w*Ri in *D. simulans, w*Ha in *D. simulans, w*Mau in *D. mauritiana, w*Sh in *D. sechellia*, and *w*Tei in *D. teissieri*. In contrast to past reports (Arnold et al. 2019; Truitt et al. 2019), we found no evidence for *w*MelCS or *w*Mel effects on *D. melanogaster T*_*p*_, which may be due to host background effects (see below). Temperature is considered a major ecological factor limiting the distribution of *Drosophila* (Hoffmann et al. 2002; Umina et al. 2005; Kellermann et al. 2009, 2012; Hoffmann 2010; Adrion et al. 2015; Rajpurohit and Schmidt 2016) and many other species (Crisp et al. 2009; Tittensor et al. 2010; Quintero and Wiens 2013). Body temperature is an important determinant of performance and fitness (Siddiqui and Barlow 1972; Huey and Berrigan 2001; Chown and Nicolson 2004; Martin and Huey 2008; Angilletta 2009; Cooper et al. 2012; Huey et al. 2012; Hoekstra et al. 2013; Condon et al. 2014), and ectotherms depend on thermoregulatory behavior to maintain body temperature within a narrow range (Angilletta et al. 2004; Martin and Huey 2008; Dillon et al. 2009; Garrity et al. 2010; Hoffmann and Sgro 2011; Rajpurohit and Schmidt 2016). Given that *Wolbachia* have spread through most insect species and other ectotherms (Werren et al. 2008; Zug and Hammerstein 2012; Weinert et al. 2015), our results motivate additional analyses of *Wolbachia* effects on *T*_*p*_ and thermoregulation of other host taxa.

Interestingly, hosts infected with A-group *Wolbachia* generally preferred cooler temperatures, whereas *D. mauritiana* infected with the B-group *w*Mau preferred warmer temperatures. Nonetheless, we did not find sufficient evidence to demonstrate these *Wolbachia* effects exhibit phylogenetic signal (Figure 2). The fitted *λ*= 0.778 was not significant (*P* = 0.203), but our simulations suggest this may be due to a small phylogeny (Supplemental Figure S5). Analyses of additional *Wolbachia*, and specifically B-group strains, are needed to test whether A- and B-group *Wolbachia* generally differ in how they influence host *T*_*p*_. Unfortunately, the only other B-group strains that infect hosts in the *D. melanogaster* subgroup (*w*No and *w*Sn) almost always occur as co-infections with other *Wolbachia* (Meany et al. 2019). *w*No co-occurs with *w*Ha in *D. simulans* (O’Neill and Karr 1990; Mercot et al. 1995; Rousset and Solignac 1995; James et al. 2002), and *w*Sn co-occurs with *w*Sh in *D. sechellia* (Giordano et al. 1995; Rousset and Solignac 1995). Naturally sampled genotypes that are singly infected with these strains are required to further assess B-group *Wolbachia* effects on host *T*_*p*_ in this subgroup. Our simulations suggest that a relatively large number of strains (*N* = 50) are required to rigorously test whether *Wolbachia* effects on *T*_*p*_ exhibit phylogenetic signal (Supplemental Figure S5). Unfortunately, measuring host *T*_*p*_ for this many *Wolbachia*-infected and -uninfected genotypes would be challenging.

The phylogenetic analysis also showed that, in some instances, closely related *Wolbachia* strains may have different effects. For example, *w*Tei and *w*Yak diverged only about 1,500 years ago and share very high sequence similarity (0.0039% third-position pairwise differences; Cooper et al. 2019), yet *w*Tei altered *T*_*p*_ of *D. teissieri* and *w*Yak had no effect on *D. yakuba* (Figure 2). Similarly, *w*Ha and *w*Sh have high sequence similarity according to our analysis (0.00008% third-position pairwise differences) and likely spread recently via introgression (Ballard 2000; Meany et al. 2019), yet our mean estimates of titer for *w*Ha in *D. simulans* (157.1) and *w*Sh in *D. sechellia* (0.2) differ by nearly three orders of magnitude (Figure 3). Host background effects may explain why closely related *Wolbachia* can have variable effects on their hosts. In a similar manner, host genomes can modify *Wolbachia* titer (Funkhouser-Jones et al. 2018), maternal *Wolbachia* transmission (Serbus and Sullivan 2007), components of host fitness (Fry et al. 2004; Dean 2006; Gruntenko et al. 2019), and the strength of cytoplasmic incompatibility (Reynolds and Hoffmann 2002; Cooper et al. 2017).

We predict that host backgrounds could explain why we found no effects of *Wolbachia* on *D. melanogaster T*_*p*_, in contrast to past reports (Arnold et al. 2019; Truitt et al. 2019). Arnold et al. (2019) found a small, yet statistically significant, reduction in *T*_*p*_ of *w*MelCS-infected *D. melanogaster* (25.06°C vs. 25.78°C for uninfected flies), and Truitt and colleagues (2019) found that a *w*MelCS variant identical to our own (according to 720 genes totaling 733,923 bp) reduced

*D. melanogaster T*_*p*_ by nearly 4°C. The effect size reported by Truitt et al. (2019) is more than two and a half times greater than the largest effect we document here for any strain, and more than five times larger than the reduction in *T*_*p*_ observed by Arnold and colleagues (2019). The *w*MelCS variant assayed in Truitt et al. (2019) was introduced into the foreign DrosDel *w*^*1118*^ isogenic background using chromosome replacement (Chrostek et al. 2013), while Arnold et al. (2019) used a standard *Oregon RC* line that was orginally established in the 1920s (Hartl and Jungen 1979; Riegler et al. 2005; Hedges et al. 2008). Our *w*MelCS-infected genotype is a sub-strain of the *Canton Special* line that was also established in the 1920s (Stern 1943; Stern and Schaeffer 1943), and sub-strains of *Canton Special* can exhibit phenotypic variation due to founder effects and drift (Colomb and Brembs 2014). Future analyses should use contemporary isofemale lines to generate reciprocally introgressed host and *Wolbachia* genotypes and dissect the relative contributions of host and *Wolbachia* genomes to *T*_*p*_, titer, and other traits.

Our temperature shift experiments indicate that changes to *T*_*p*_ of infected host genotypes generally do not alter *Wolbachia* titer, but in a few instances, reductions in *T*_*p*_ seem to increase *Wolbachia* replication within host bodies (Figure 3). *w*Sh-infected *D. sechellia* and *w*Tei-infected *D. teissieri* preferred cooler temperatures than uninfected flies (Figure 2), and infected males reared at 21.5°C had significantly higher *Wolbachia* titer when shifted to a cold 18°C treatment for 24 hours, compared to a warm 25°C treatment (Figure 3). Moghadam et al. (2018) reported a similar effect of cold temperature on *Wolbachia* titer in male *D. melanogaster*, in which males developed at 13°C had higher microbial diversity and a higher relative abundance of *Wolbachia* than males developed at 23° and 31°C (based on 16S rRNA sequencing). Our results are consistent with a hypothesis of parasite manipulation, in which *Wolbachia* alter host behavior to seek environmental conditions that promote *Wolbachia* growth and transmission (Poulin 2010; De Roode and Lefèvre 2012; Curtis 2014; Heil 2016; Vale et al. 2018; Weinersmith 2019). However, we found no temperature-associated increases in titer for *w*Sh- and *w*Tei-infected females or for any other *Wolbachia* strains we assessed. Other studies have also reported male-biased environmental effects on *Wolbachia* titer (Correa and Ballard 2012; Foo et al. 2019; Hague et al. 2020); for example, our own work demonstrated that maternal transmission of *w*Yak to sons is more efficient than to daughters when *D. yakuba* mothers are reared in cold 20°C conditions (Hague et al. 2020).

Our findings do not provide support for the hypothesis proposed by Truitt et al. (2019) that modifications to *T*_*p*_ represent an adaptive host response (e.g., behavioral chill) to reduce *Wolbachia* titer and mitigate the negative effects of infection. In particular, Truitt et al. (2019) speculated that *w*MelCS is costly to the host because the strain has a higher titer and growth rate than *w*Mel (Chrostek et al. 2013), and *w*MelCS-infected *D. melanogaster* prefer colder temperatures to reduce *Wolbachia* titer and limit costly infections. The authors did not measure *w*MelCS titer or estimate host fitness components to test this hypothesis (Truitt et al. 2019). We found no effects of *w*MelCS or *w*Mel on *T*_*p*_ of *D. melanogaster* and no evidence that decreases in *T*_*p*_ reduce *Wolbachia* titer for other infected systems (Figure 3). Nonetheless, the observation that most *Wolbachia*-infected hosts have altered *T*_*p*_ motivates future analyses of host behaviors that might mitigate negative aspects of infection, especially because *Wolbachia* can have costly effects on hosts (Hoffmann et al. 1990; Stouthamer and Luck 1993; Turelli and Hoffmann 1995; Weeks et al. 2007). We found no association between changes to *T*_*p*_ and a decrease in adult *Wolbachia* titer, but perhaps infected females seek oviposition sites that reduce the efficiency of *Wolbachia* maternal transmission (Dillon et al. 2009). *Wolbachia* maternal transmission seems to be influenced by relatively cold 20°C temperatures in *Drosophila* (Hague et al. 2020) and hot temperatures in mosquitoes (Ross et al. 2017, 2019a). Future work should test whether rearing infected hosts at their *T*_*p*_ influences *Wolbachia* titer.

Our results add to mounting literature showing that temperature is an important abiotic factor mediating interactions between *Wolbachia* and their hosts (Charlesworth et al. 2019). In particular, *Wolbachia* titer seems to be especially sensitive to temperature (Clancy and Hoffmann 1998; Mouton et al. 2006, 2007; Bordenstein and Bordenstein 2011; Ross et al. 2017, 2019a; Sumi et al. 2017; Hague et al. 2020). Our 24-hour temperature shift experiments suggest *Wolbachia* titer can change over very short time periods due to environmental conditions. Mouton et al. (2006) reported a similar result in which *Leptopilina heterotoma* infected with *w*Lhet experienced a significant increase in *Wolbachia* titer over the course of a single host generation when mothers were reared at 20°C and daughters developed at 26°C. Temperature-induced changes to *Wolbachia* titer are likely to have cascading effects, given that titer influences other host phenotypes (López-Madrigal and Duarte 2019). For example, exposure to heat stress is associated with correlated declines in *Wolbachia* titer and the severity of cytoplasmic incompatibility in *w*Mel-transinfected *Ae. aegypti* (Ross et al. 2017, 2019a). In *Drosophila* hosts, temperature has been shown to modify the strength of cytoplasmic incompatibility (Hoffmann et al. 1986, 1990; Clancy and Hoffmann 1998; Reynolds and Hoffmann 2002), maternal transmission (Turelli and Hoffmann 1995; Hague et al. 2020), and host fitness effects (Olsen et al. 2001; Versace et al. 2014; Kriesner et al. 2016). Clearly, more work on how temperature influences *Wolbachia*-host interactions is needed.

## Conclusion

We show that A- and B-group *Wolbachia* induce changes to host *T*_*p*_, and short 24-hour shifts in temperature can increase titer in some *Wolbachia*-infected males. Behavioral changes like these are likely to have fundamental consequences for host physiology and thermoregulation. *Wolbachia* also modify a range of other ecologically important host traits in *Drosophila* species, including reproduction (Hoffmann and Turelli 1997; Werren et al. 2008), virus blocking (Hedges et al. 2008; Teixeira et al. 2008; Osborne et al. 2009; Martinez et al. 2014), nutrient provisioning (Brownlie et al. 2009; Newton and Rice 2020), and activity levels (van Houte et al. 2013; Bi and Wang 2019). Given that *T*_*p*_ and many other *Drosophila* traits vary clinally (Hoffmann and Weeks 2007; Rajpurohit and Schmidt 2016), future studies should consider the role of *Wolbachia* in classic *Drosophila* clines (Adrion et al. 2015). For example, *w*Mel infection frequencies (Kriesner et al. 2016) and the *T*_*p*_ of *D. melanogaster* (Rajpurohit and Schmidt 2016) both vary spatially in eastern North America.

Understanding the impact of *Wolbachia* on host performance and fitness is crucial for predicting evolutionary outcomes of *Wolbachia*-host interactions (Ross et al. 2019b). The initial spread of *Wolbachia* through new host populations is driven by beneficial effects on host fitness that cause infections to deterministically spread from low initial frequencies (Caspari and Watson 1959; Hoffmann et al. 1990; Barton and Turelli 2011). Yet, strong positive host effects have not been directly connected to spread in nature for any *Wolbachia*-infected host species (Cooper et al. 2017; Shi et al. 2018; Meany et al. 2019; Ross et al. 2019b); although, *w*Ri recently evolved to confer a 10% fecundity advantage to *D. simulans* (Weeks et al. 2007). Few data exist for other components of host fitness, but protection from viruses and nutrient provisioning remain candidates for potential host benefits (Hedges et al. 2008; Teixeira et al. 2008; Brownlie et al. 2009; Osborne et al. 2009; Martinez et al. 2014; Nikoh et al. 2014; Newton and Rice 2019; but see Shi et al. 2018). Basic research on how *Wolbachia* modify different components of host fitness, like the effects on *T*_*p*_ reported here, represents a key step to uncovering how *Wolbachia* benefit hosts and spread to become a global pandemic.

## ACKNOWLEDGEMENTS

We thank Tim Wheeler for assistance in the lab and Will Conner for help with bioinformatic analyses. Isaac Humble helped construct the thermal gradient apparatus. Dave Begun, Michael Turelli, and Daniel Matute kindly provided flies used in this study. The Cooper lab group, Michael May, and Gregg Thomas provided valuable feedback that improved the quality of the manuscript. We thank the Genomics Core and the Environmental Control for Organismal Research (ECOR) Laboratories at the University of Montana for their support.

## SUPPLEMENTAL MATERIALS

**Supplemental Figure S1.**
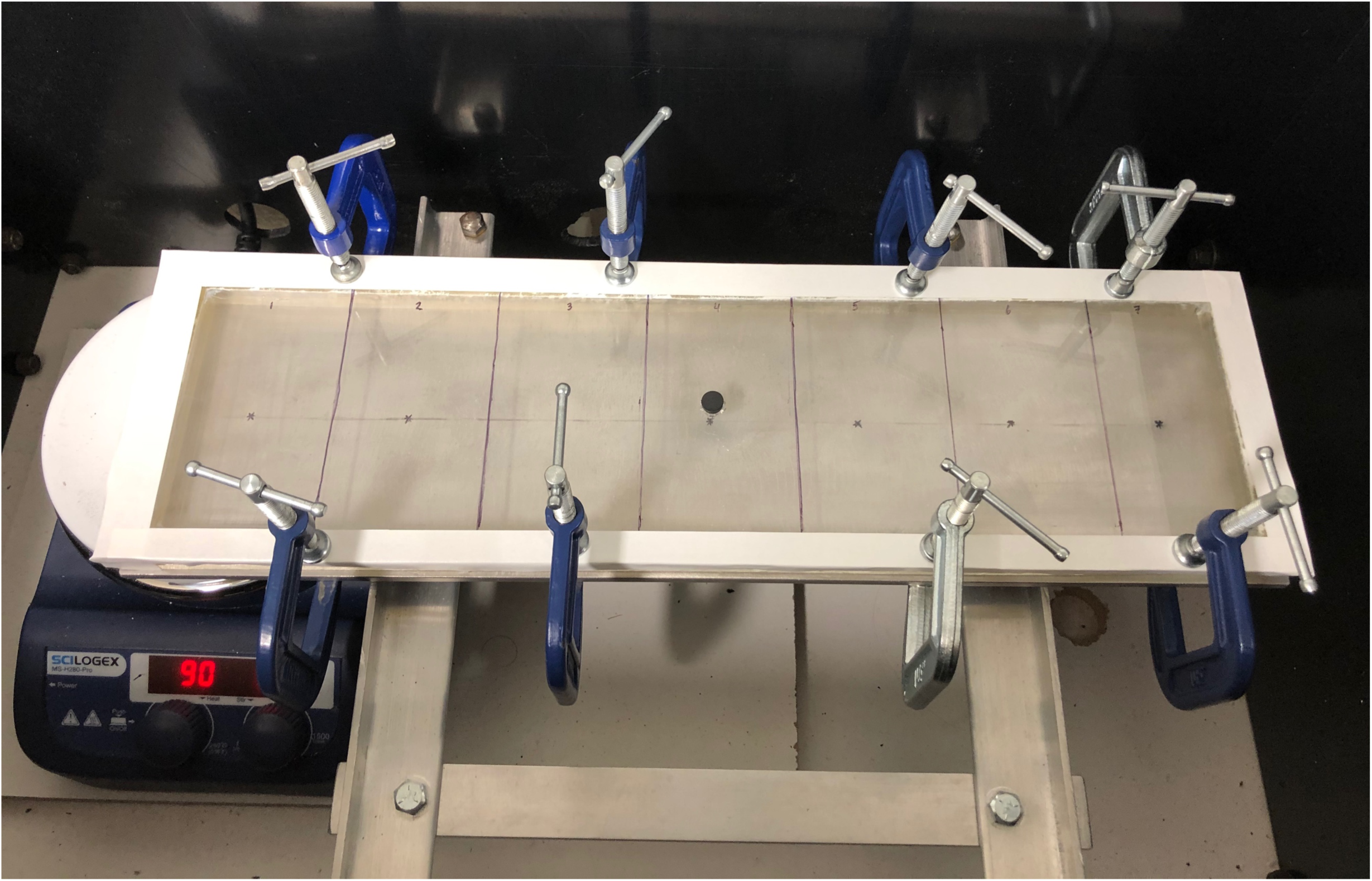
The thermal gradient apparatus is composed of a 44 x 13 x 1 cm aluminum plate and a 1 cm-high removable Plexiglas lid. The thermal gradient is subdivided into seven 10 x 6 cm sections (Supplemental Figure S2, Table S2).

**Supplemental Figure S2.**
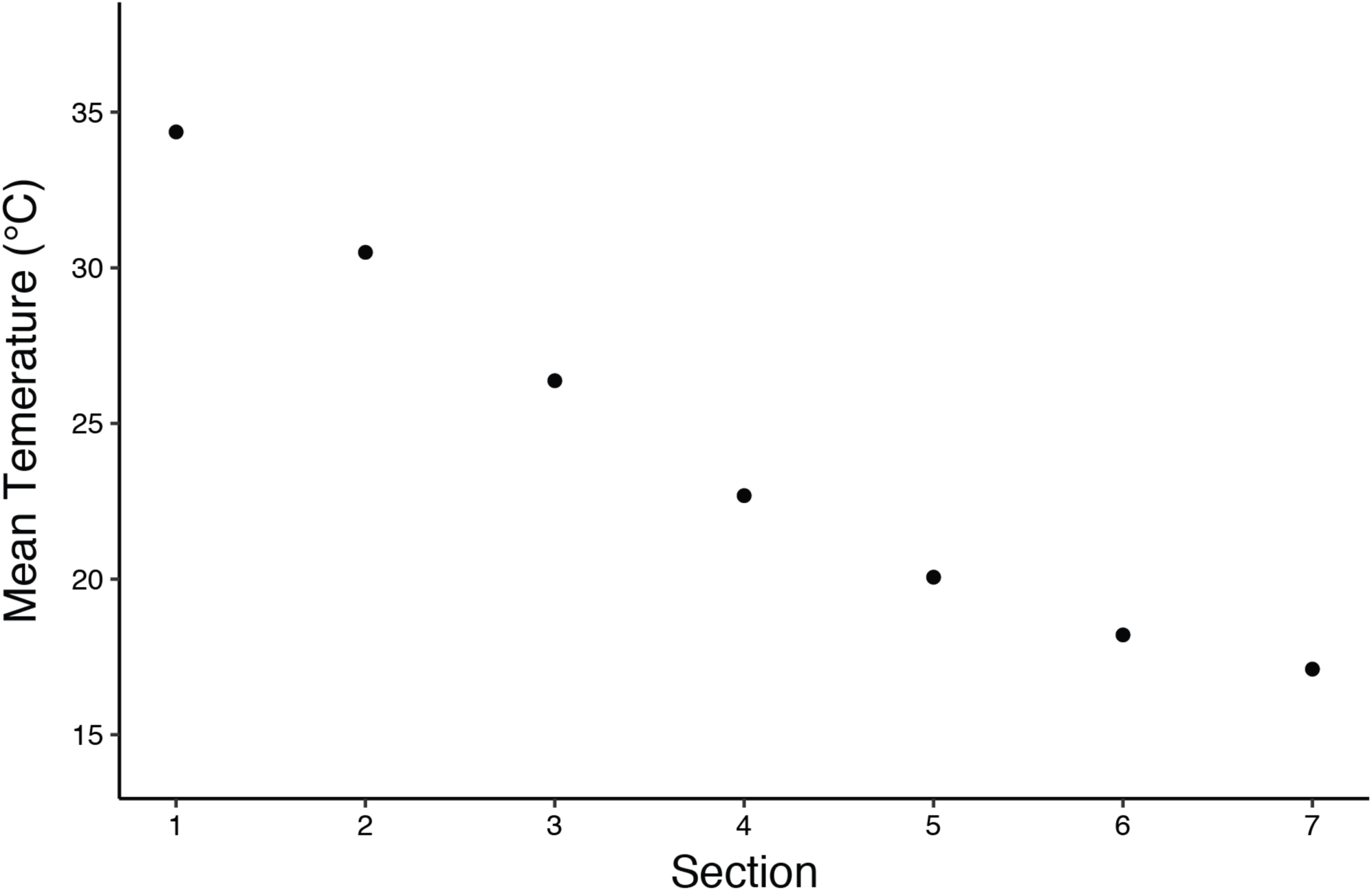
Mean temperature of each section on the thermal gradient apparatus. Mean and standard errors are presented in Supplemental Table S2.

**Supplemental Figure S3.**
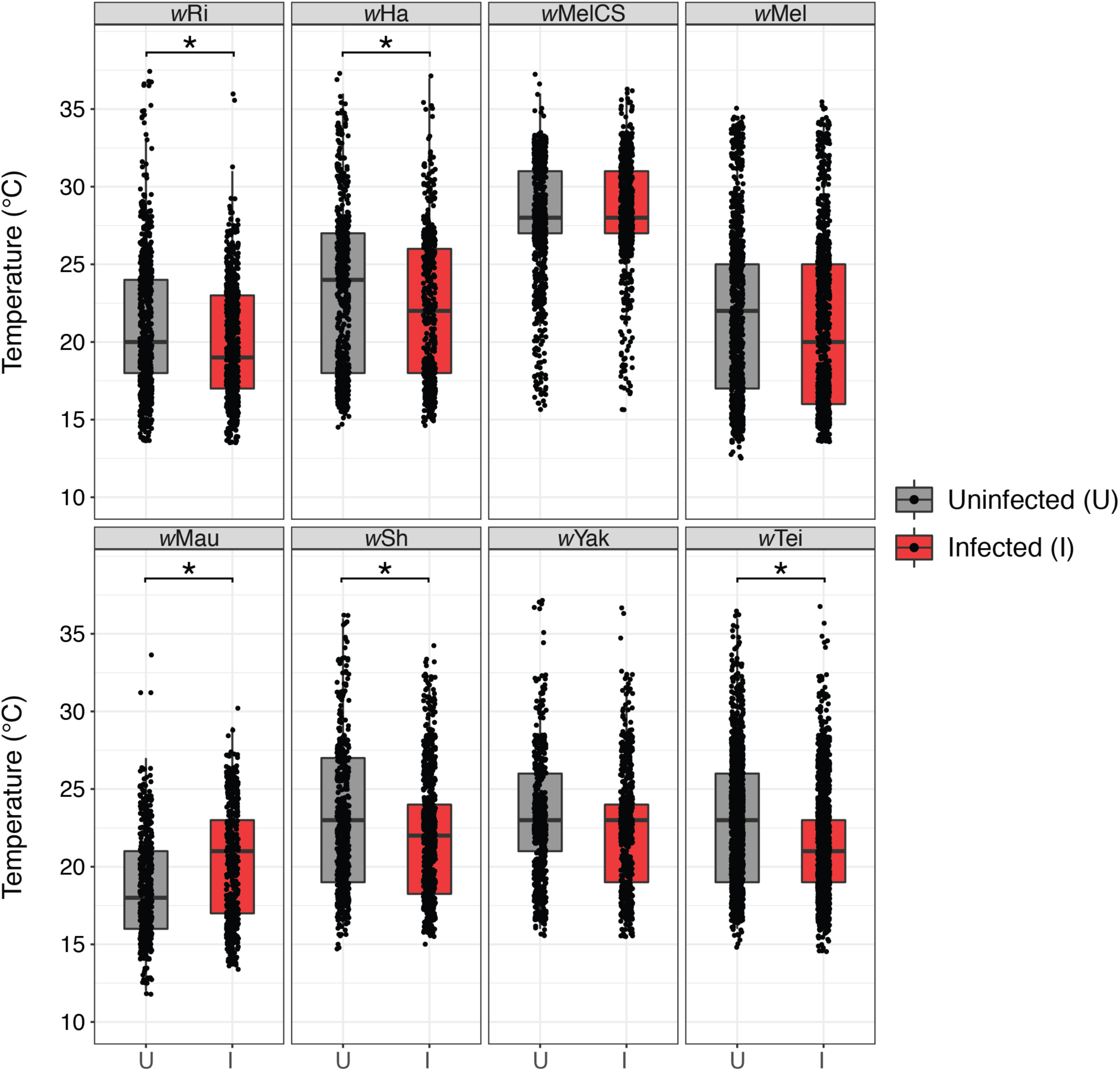
Box plots showing temperature preference (*T*_*p*_) for uninfected (U) and infected (I) flies of each genotype when the coldest section of the thermal gradient (section 7) is included. Asterisks denote a significant main effect of *Wolbachia* infection on *T*_*p*_ from the GLMMs (Supplemental Table S4). Individual points are jittered to show overlap.

**Supplemental Figure S4.**
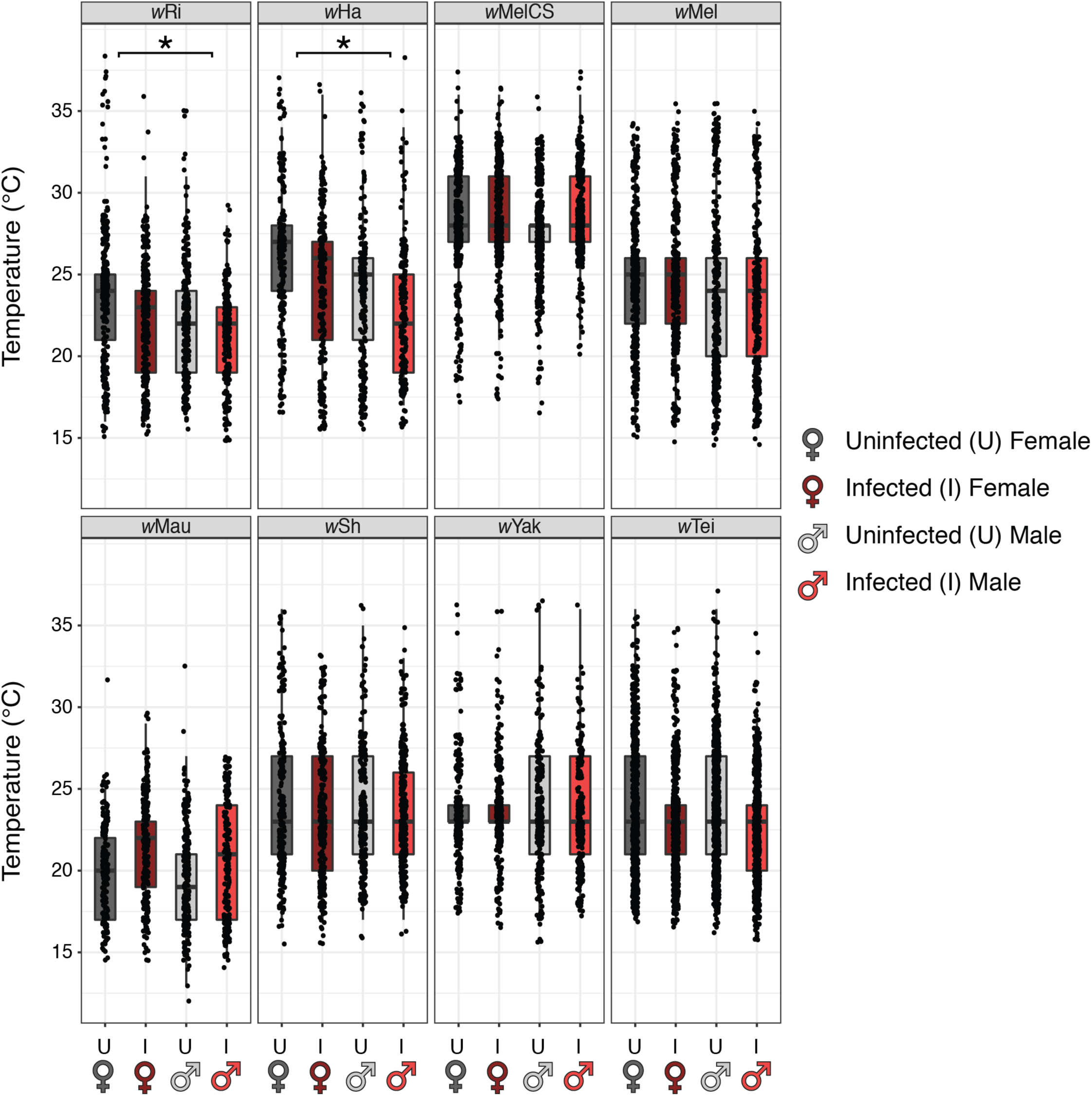
Box plots showing temperature preference (*T*_*p*_) for uninfected (U) and infected (I) flies of each genotype, separated by sex. Asterisks denote a significant main effect of sex on *T*_*p*_ from the GLMMs (Table 1). Individual points are jittered to show overlap. The coldest section of the thermal gradient apparatus (section 7) is removed from the dataset.

**Supplemental Figure S5.**
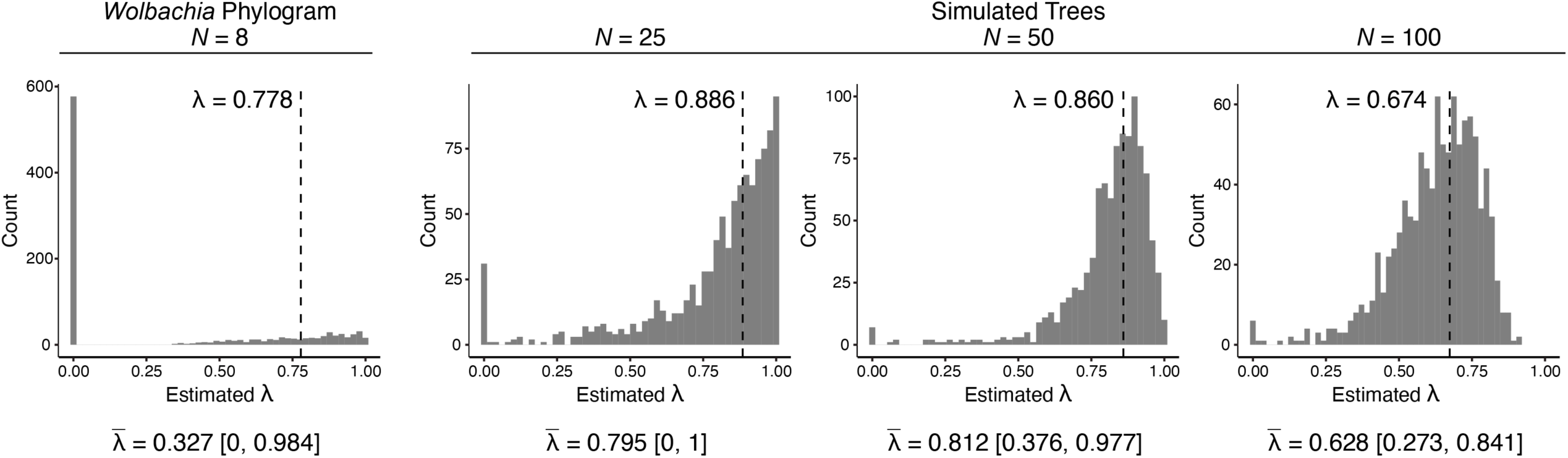
Distribution of maximum likelihood estimates of *λ* from 1,000 bootstrap replicates. The bootstrap analysis for our *Wolbachia* phylogram (Figure 2) is show to the left. To the right are simulated phylogenies with an increasing number of *Wolbachia* strains included (*N* = 25, 50, 100). For simulated trees, character evolution was simulated with our *λ* estimate of 0.778 using the “sim.bdtree” and “sim.char” functions in the *geiger* R package (Harmon et al. 2008). For each graph, fitted *λ* values for the original phylogeny are shown above with a vertical dashed line. Note that fitted *λ* values for the simulated phylogenies differ slightly from *λ* = 0.778, because “sim.char” uses a Brownian-motion model to simulate character evolution along the phylogeny. Below each graph, the mean estimate of *λ* from the 1,000 replicates 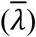 is shown with associated 95% confidence intervals. The bootstrapping analyses generally show that small phylogenies (*N* = 8, 25) have a large number of near-zero *λ* values arising by random chance, which increases the uncertainty of parameter estimation. As the number of strains increases (*N* = 50, 100), bootstrapped estimates of *λ* cluster around the true *λ* value fitted to the original phylogeny.

**Supplemental Table S1.** Genotype IDs for different *Wolbachia*-infected host species used in this study.

| Species | *Wolbachia* | Genotype ID |
| --- | --- | --- |
| *D. simulans* | wRi | *Riv84* |
| *D. simulans* | wHa | *Car5* |
| *D. melanogaster* | wMelCS | *Canton S Berkeley* |
| *D. melanogaster* | wMel | *PC75* |
| *D. mauritiana* | wMau | *mauR31* |
| *D. sechellia* | wSh | *PmuseumbananaI* |
| *D. yakuba* | wYak | *B13L5* |
| *D. teissieri* | wTei | *B13L11* |

**Supplemental Table S2.** Mean temperature and standard error for each section of the custom-built thermal gradient apparatus, across all 379 experimental replicates in this study.

| Section | Mean Temp. (°C) | Std. Error |
| --- | --- | --- |
| 1 | 34.4 | 0.08 |
| 2 | 30.5 | 0.08 |
| 3 | 26.4 | 0.07 |
| 4 | 22.7 | 0.06 |
| 5 | 20.1 | 0.07 |
| 6 | 18.2 | 0.06 |
| 7 | 17.1 | 0.06 |

**Supplemental Table S3.** *T*_*p*_ data for the *w*Ha, *w*MelCS, *w*Mel, and *w*Mau host genotypes approximated a normal distribution. Results from LMMs for each *T*_*p*_ dataset are shown below. Statistically significant fixed effects at *P* < 0.05 are marked in bold text with asterisks.

| | $w$Ri | | | $w$Ha | | | $w$MelCS | | | $w$Mel | | |
| --- | --- | --- | --- | --- | --- | --- | --- | --- | --- | --- | --- | --- |
| Explanatory variable | coefficient | $\chi^2$ | *P* value | coefficient | $\chi^2$ | *P* value | coefficient | $\chi^2$ | *P* value | coefficient | $\chi^2$ | *P* value |
| Infection Status | 1.607 | 5.082 | **0.024\*** | 1.576 | 5.201 | **0.023\*** | -0.492 | 1.065 | 0.302 | -0.101 | 0.027 | 0.871 |
| Sex | -1.360 | 3.544 | 0.060 | -1.640 | 5.438 | **0.020\*** | -0.197 | 0.155 | 0.694 | -1.125 | 3.178 | 0.075 |
| Age | -0.160 | 0.092 | 0.761 | -0.048 | 0.058 | 0.810 | -0.574 | 9.683 | **0.002\*** | 0.316 | 2.165 | 0.141 |
| Run Order | 0.026 | 0.013 | 0.909 | 0.167 | 0.533 | 0.465 | 0.314 | 3.613 | 0.057 | 0.103 | 0.264 | 0.608 |
| Infection * Sex | -0.383 | 0.141 | 0.707 | -0.457 | 0.206 | 0.650 | 0.023 | 0.001 | 0.974 | 0.528 | 0.357 | 0.550 |
| Sample Size | 1015 |  |  | 857 |  |  | 1727 |  |  | 1341 |  |  |

| | $w$Mau | | | $w$Sh | | | $w$Yak | | | $w$Tei | | |
| --- | --- | --- | --- | --- | --- | --- | --- | --- | --- | --- | --- | --- |
| Explanatory variable | coefficient | $\chi^2$ | *P* value | coefficient | $\chi^2$ | *P* value | coefficient | $\chi^2$ | *P* value | coefficient | $\chi^2$ | *P* value |
| Infection Status | -1.871 | 5.086 | **0.024\*** | 1.019 | 4.336 | **0.037\*** | 0.035 | 0.003 | 0.958 | 0.734 | 5.573 | **0.018\*** |
| Sex | -1.088 | 2.127 | 0.145 | 0.261 | 0.314 | 0.575 | -0.248 | 0.143 | 0.705 | -0.469 | 2.300 | 0.129 |
| Age | -0.349 | 1.509 | 0.219 | 0.072 | 0.109 | 0.742 | 0.608 | 2.814 | 0.093 | -0.257 | 2.305 | 0.129 |
| Run Order | 0.456 | 3.065 | 0.080 | 0.253 | 2.673 | 0.102 | 0.223 | 1.073 | 0.300 | 0.022 | 0.049 | 0.824 |
| Infection * Sex | 0.793 | 0.507 | 0.477 | -0.799 | 1.386 | 0.239 | 0.291 | 0.097 | 0.755 | 0.421 | 0.936 | 0.333 |
| Sample Size | 818 |  |  | 1087 |  |  | 1056 |  |  | 2500 |  |  |

**Supplemental Table S4.** Results and sample sizes from the GLMMs and LMMs including the flies located in the coldest section of the thermal gradient apparatus (section 7). Statistically significant fixed effects at *P* < 0.05 are marked in bold text with asterisks.

GLMMs
| Explanatory variable | wRi |  |  | wHa |  |  | wMelCS |  |  | wMel |  |  |
| --- | --- | --- | --- | --- | --- | --- | --- | --- | --- | --- | --- | --- |
| | coefficient | $\chi^2$ | *P* value | coefficient | $\chi^2$ | *P* value | coefficient | $\chi^2$ | *P* value | coefficient | $\chi^2$ | *P* value |
| Infection Status | 0.063 | 4.757 | **0.029\*** | 0.066 | 6.513 | **0.011\*** | -0.022 | 1.463 | 0.226 | 0.036 | 1.788 | 0.181 |
| Sex | -0.066 | 5.115 | **0.024\*** | -0.060 | 5.057 | **0.025\*** | -0.007 | 0.156 | 0.693 | -0.045 | 2.743 | 0.098 |
| Age | 0.001 | 0.002 | 0.968 | 0.005 | 0.394 | 0.530 | -0.024 | 12.093 | **0.001\*** | 0.008 | 0.640 | 0.424 |
| Run Order | 0.001 | 0.011 | 0.916 | 0.009 | 1.056 | 0.304 | 0.015 | 5.733 | **0.017\*** | 0.020 | 5.374 | **0.020\*** |
| Infection * Sex | -0.028 | 0.454 | 0.501 | -0.020 | 0.281 | 0.596 | 0.003 | 0.016 | 0.900 | 0.012 | 0.099 | 0.753 |
| Sample Size | 1534 |  |  | 1135 |  |  | 1770 |  |  | 1962 |  |  |

| Explanatory variable | wMau |  |  | wSh |  |  | wYak |  |  | wTei |  |  |
| --- | --- | --- | --- | --- | --- | --- | --- | --- | --- | --- | --- | --- |
| | coefficient | $\chi^2$ | *P* value | coefficient | $\chi^2$ | *P* value | coefficient | $\chi^2$ | *P* value | coefficient | $\chi^2$ | *P* value |
| Infection Status | -0.110 | 7.615 | **0.006\*** | 0.047 | 5.835 | **0.016\*** | 0.012 | 0.270 | 0.603 | 0.045 | 9.814 | **0.002\*** |
| Sex | -0.052 | 2.055 | 0.152 | 0.018 | 0.962 | 0.327 | -0.028 | 1.450 | 0.229 | -0.019 | 1.796 | 0.180 |
| Age | -0.015 | 1.116 | 0.291 | 0.002 | 0.078 | 0.780 | 0.028 | 4.695 | **0.030\*** | -0.013 | 3.039 | 0.081 |
| Run Order | 0.022 | 3.045 | 0.081 | 0.018 | 8.319 | **0.004\*** | 0.010 | 1.874 | 0.171 | 0.004 | 0.941 | 0.332 |
| Infection * Sex | 0.063 | 1.383 | 0.240 | -0.026 | 0.920 | 0.337 | 0.022 | 0.450 | 0.502 | 0.018 | 0.808 | 0.369 |
| Sample Size | 1009 |  |  | 1351 |  |  | 1232 |  |  | 2951 |  |  |

LMMs
| Explanatory variable | wRi |  |  | wHa |  |  | wMelCS |  |  | wMel |  |  |
| --- | --- | --- | --- | --- | --- | --- | --- | --- | --- | --- | --- | --- |
| | coefficient | $\chi^2$ | *P* value | coefficient | $\chi^2$ | *P* value | coefficient | $\chi^2$ | *P* value | coefficient | $\chi^2$ | *P* value |
| Infection Status | 1.356 | 4.142 | **0.042\*** | 1.548 | 5.477 | **0.019\*** | -0.604 | 1.247 | 0.264 | 0.816 | 1.736 | 0.188 |
| Sex | -1.344 | 4.077 | **0.043\*** | -1.315 | 3.864 | **0.049\*** | -0.189 | 0.109 | 0.741 | -0.942 | 2.307 | 0.129 |
| Age | 0.052 | 0.011 | 0.915 | 0.112 | 0.341 | 0.559 | -0.671 | 10.257 | **0.001\*** | 0.174 | 0.657 | 0.418 |
| Run Order | 0.021 | 0.010 | 0.922 | 0.194 | 0.792 | 0.373 | 0.399 | 4.544 | **0.033\*** | 0.435 | 4.733 | **0.030\*** |
| Infection * Sex | -0.655 | 0.483 | 0.487 | -0.541 | 0.318 | 0.573 | 0.094 | 0.014 | 0.907 | 0.213 | 0.059 | 0.809 |
| Sample Size | 1534 |  |  | 1135 |  |  | 1770 |  |  | 1962 |  |  |

| Explanatory variable | wMau |  |  | wSh |  |  | wYak |  |  | wTei |  |  |
| --- | --- | --- | --- | --- | --- | --- | --- | --- | --- | --- | --- | --- |
| | coefficient | $\chi^2$ | *P* value | coefficient | $\chi^2$ | *P* value | coefficient | $\chi^2$ | *P* value | coefficient | $\chi^2$ | *P* value |
| Infection Status | -2.232 | 5.971 | **0.015\*** | 1.030 | 4.614 | **0.032\*** | 0.286 | 0.218 | 0.641 | 0.907 | 7.640 | **0.006\*** |
| Sex | -1.048 | 1.611 | 0.204 | 0.392 | 0.732 | 0.392 | -0.634 | 1.112 | 0.292 | -0.502 | 2.416 | 0.120 |
| Age | -0.288 | 0.836 | 0.360 | 0.055 | 0.067 | 0.796 | 0.639 | 3.658 | 0.056 | -0.280 | 2.475 | 0.116 |
| Run Order | 0.441 | 2.326 | 0.127 | 0.397 | 6.853 | **0.009\*** | 0.249 | 1.581 | 0.209 | 0.100 | 0.943 | 0.331 |
| Infection * Sex | 1.292 | 1.102 | 0.294 | -0.624 | 0.875 | 0.350 | 0.486 | 0.319 | 0.572 | 0.515 | 1.257 | 0.262 |
| Sample Size | 1009 |  |  | 1351 |  |  | 1232 |  |  | 2951 |  |  |

**Supplemental Table S5.** The scaffold count, N50, and total assembly size of each *Wolbachia* assembly.

| Genome | Host Genotype ID | Scaffold Count | N50 | Total Assembly Size |
| --- | --- | --- | --- | --- |
| wMel | PC75 | 76 | 29,986 | 1,237,996 |
| wMelCS | Canton S Berkeley | 89 | 25,105 | 1,233,787 |
| wSh | LD15 (Accession SRX3029362) | 83 | 25,919 | 1,294,885 |

**Supplemental Table S6.**
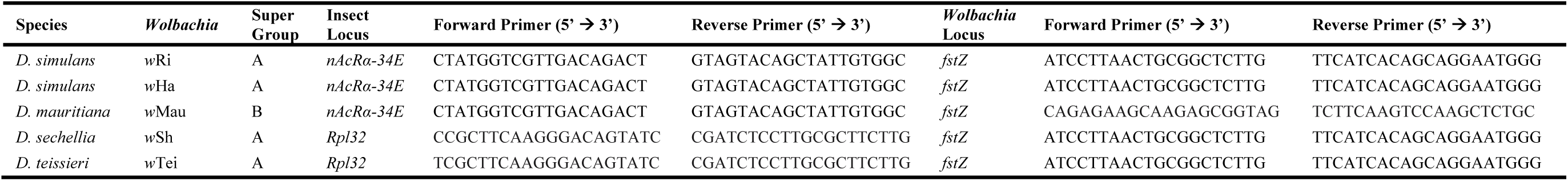
qPCR primers used to measure relative *Wolbachia* density in temperature shift experiments.

**Supplemental Table S7.** Results from *T*_*p*_ assays. Sample sizes (n) and results are shown for uninfected and infected flies of each genotype. Results for uninfected and infected flies are also separated by sex.

| Species | *Wolbachia* | Infection Status | n | Mean $T_p$ (°C) | Standard Error |
| --- | --- | --- | --- | --- | --- |
| *D. simulans* | wRi | Uninfected | 504 | 23.1 | 0.18 |
|  |  | Female | 272 | 23.9 | 0.25 |
|  |  | Male | 232 | 22.2 | 0.24 |
|  |  | Infected | 511 | 21.7 | 0.14 |
|  |  | Female | 280 | 22.3 | 0.21 |
|  |  | Male | 231 | 21.1 | 0.17 |
| *D. simulans* | wHa | Uninfected | 458 | 25.0 | 0.21 |
|  |  | Female | 239 | 25.9 | 0.27 |
|  |  | Male | 219 | 23.9 | 0.30 |
|  |  | Infected | 399 | 23.7 | 0.22 |
|  |  | Female | 218 | 24.5 | 0.30 |
|  |  | Male | 181 | 22.7 | 0.31 |
| *D. melanogaster* | wMelCS | Uninfected | 822 | 27.9 | 0.11 |
|  |  | Female | 376 | 28.0 | 0.17 |
|  |  | Male | 446 | 27.8 | 0.14 |
|  |  | Infected | 905 | 28.4 | 0.09 |
|  |  | Female | 452 | 28.2 | 0.14 |
|  |  | Male | 453 | 28.5 | 0.11 |
| *D. melanogaster* | wMel | Uninfected | 702 | 24.3 | 0.18 |
|  |  | Female | 398 | 24.6 | 0.20 |
|  |  | Male | 304 | 24.0 | 0.30 |
|  |  | Infected | 639 | 24.1 | 0.18 |
|  |  | Female | 337 | 24.6 | 0.23 |
|  |  | Male | 302 | 23.6 | 0.28 |
| *D. mauritiana* | wMau | Uninfected | 405 | 19.8 | 0.15 |
|  |  | Female | 177 | 20.2 | 0.22 |
|  |  | Male | 228 | 19.5 | 0.21 |
|  |  | Infected | 413 | 21.1 | 0.16 |
|  |  | Female | 200 | 21.5 | 0.22 |
|  |  | Male | 213 | 20.7 | 0.24 |
| *D. sechellia* | wSh | Uninfected | 501 | 23.9 | 0.17 |
|  |  | Female | 233 | 24.1 | 0.27 |
|  |  | Male | 268 | 23.8 | 0.22 |
|  |  | Infected | 586 | 23.4 | 0.15 |
|  |  | Female | 279 | 23.3 | 0.24 |
|  |  | Male | 307 | 23.5 | 0.18 |
| *D. yakuba* | wYak | Uninfected | 536 | 23.7 | 0.15 |
|  |  | Female | 277 | 23.6 | 0.19 |
|  |  | Male | 259 | 23.8 | 0.23 |
|  |  | Infected | 520 | 23.6 | 0.14 |
|  |  | Female | 273 | 23.7 | 0.19 |
|  |  | Male | 247 | 23.5 | 0.21 |
| *D. teissieri* | wTei | Uninfected | 1337 | 23.7 | 0.10 |
|  |  | Female | 627 | 23.8 | 0.16 |
|  |  | Male | 710 | 23.6 | 0.13 |
|  |  | Infected | 1163 | 22.7 | 0.10 |
|  |  | Female | 552 | 22.9 | 0.14 |
|  |  | Male | 611 | 22.5 | 0.13 |

**Supplemental Table S8.** Titer results from 24-hour temperature shift experiments.

| Species | *Wolbachia* | Temperature | Sex | Biological Replicates | Median Relative *Wolbachia* Density ($2^{\Delta Ct}$) | Lower Quartile | Upper Quartile |
| --- | --- | --- | --- | --- | --- | --- | --- |
| *D. simulans* | wRi | 18°C | F | 5 | 385.34 | 240.52 | 617.37 |
|  |  | 18°C | M | 6 | 419.90 | 349.05 | 531.99 |
|  |  | 25°C | F | 5 | 350.52 | 246.14 | 604.67 |
|  |  | 25°C | M | 6 | 419.80 | 280.70 | 573.06 |
| *D. simulans* | wHa | 18°C | F | 6 | 105.56 | 83.96 | 136.76 |
|  |  | 18°C | M | 6 | 112.02 | 104.86 | 124.60 |
|  |  | 25°C | F | 6 | 125.36 | 107.02 | 156.51 |
|  |  | 25°C | M | 6 | 107.15 | 102.15 | 123.32 |
| *D. mauritiana* | wMau | 18°C | F | 6 | 30.37 | 26.07 | 43.12 |
|  |  | 18°C | M | 6 | 25.97 | 12.83 | 45.55 |
|  |  | 25°C | F | 6 | 58.07 | 24.01 | 108.83 |
|  |  | 25°C | M | 6 | 30.21 | 21.96 | 37.19 |
| *D. sechellia* | wSh | 18°C | F | 6 | 0.24 | 0.23 | 0.25 |
|  |  | 18°C | M | 6 | 0.16 | 0.14 | 0.16 |
|  |  | 25°C | F | 6 | 0.33 | 0.24 | 0.36 |
|  |  | 25°C | M | 6 | 0.11 | 0.09 | 0.13 |
| *D. teissieri* | wTei | 18°C | F | 6 | 3.56 | 3.42 | 3.73 |
|  |  | 18°C | M | 6 | 3.36 | 3.20 | 3.55 |
|  |  | 25°C | F | 6 | 3.23 | 3.15 | 3.53 |
|  |  | 25°C | M | 6 | 2.98 | 2.78 | 3.12 |

